# Overlapping long sequence reads: Current innovations and challenges in developing sensitive, specific and scalable algorithms

**DOI:** 10.1101/081596

**Authors:** Justin Chu, Hamid Mohamadi, René L Warren, Chen Yang, Inanc Birol

## Abstract

Identifying overlaps between error-prone long reads, specifically those from Oxford Nanopore Technologies (ONT) and Pacific Biosciences (PB), is essential for certain downstream applications, including error correction and *de novo* assembly. Though akin to the read-to-reference alignment problem, read-to-read overlap detection is a distinct problem that can benefit from specialized algorithms that perform efficiently and robustly on high error rate long reads. Here, we review the current state-of-the-art read-to-read overlap tools for error-prone long reads, including BLASR, DALIGNER, MHAP, GraphMap, and Minimap. These specialized bioinformatics tools differ not just in their algorithmic designs and methodology, but also in their robustness of performance on a variety of datasets, time and memory efficiency, and scalability. We highlight the algorithmic features of these tools, as well as their potential issues and biases when utilizing any particular method. We benchmarked these tools, tracking their resource needs and computational performance, and assessed the specificity and precision of each. The concepts surveyed may apply to future sequencing technologies, as scalability is becoming more relevant with increased sequencing throughput.

**Supplementary information:** Supplementary data are available at *Bioinformatics* online.

## 1 Introduction

As today’s lion share of DNA and RNA sequencing is carried out on Illumina sequencing instruments (San Diego, CA), most *de novo* assembly methods are optimized expecting short read data with an error rate less than 1% (Laehnemann, et al., 2016; Ross, et al., 2013). However, their associated short read length and GC bias bring significant challenges for downstream analyses (Ross, et al., 2013; Smith, et al., 2008). For instance, short read lengths make it difficult to assemble entire genomes due to repetitive elements (Alkan, et al., 2010). The development of paired-end and mate-pair sequencing library protocols has helped mitigate this problem (Potato Genome Sequencing Consortium, 2011), but is still a long-standing computational problem for *de novo* genome assembly (Treangen and Salzberg, 2012). Co-localization of short reads is a potential strategy to increase the contiguity of assemblies, using technologies such as Illumina TruSeq synthetic long reads (McCoy, et al., 2014) and 10X Genomics Chromium (Pleasanton, CA) (Eisenstein, 2015); however, tandem repeats in the same long single DNA fragment may continue confounding assembly methodologies. In that respect, long sequencing holds great promise, and has proved useful in resolving such issues (Ummat and Bashir, 2014). Still, the appreciable error rates associated with technologies offered by Oxford Nanopore Technologies (Oxford, UK; ONT) and Pacific Biosciences (Menlo Park, CA; PB) pose new challenges for the *de novo* assembly problem.

Read-to-read overlap detection is typically the first step of *de novo* Overlap-Layout-Consensus (OLC) assembly, which dominate the assembly of long reads (Berlin, et al., 2015; Loman, et al., 2015). The first long read *de novo* assembly methods employed error correction as their initial pipeline step (Berlin, et al., 2015; Chin, et al., 2013; Loman, et al., 2015), which also requires read-to-read overlaps. Alternatively, one can forego the error correction stages of assembly in favor of overlap between uncorrected raw reads (Li, 2016). The benefits of an uncorrected read-to-read overlap paradigm for assembly include a lower computation cost, and repressing artifacts that may arise from read correction, such as collapsed homologous regions. On the other hand, for these methods, correctness of the initial set of overlaps are even more critical. Further, overlap detection has been identified as the efficiency bottleneck when using OLC assembly methodology (Myers, 2014) for large genomes.

At present, multiple tools are capable of overlapping error-prone long read data at varying levels of accuracy. These methods differ in their methodology, but have some common aspects, such as the use of short exact subsequences (seeds) to discover candidate overlaps. Here we provide an overview of how each of these tool work, along with the conceptual motivations within their design.

## 2 Background

### 2.1 Current challenges when using PB sequencing

PB sequencing uses a DNA polymerase anchored in a well small enough to act as a zero-mode waveguide (ZMW) (Levene, et al., 2003). The polymerase acts on a single DNA molecule incorporating fluorophores labeled nucleotides, which are excited by a laser. The resulting optical signal is recorded by a high-speed camera in real time (Eid, et al., 2009). Base calling errors on this platform occur at a rate of around 16% (Laehnemann, et al., 2016), and are dominated by insertions (Carneiro, et al., 2012; Ross, et al., 2013), which are possibly caused by the dissociation of cognate nucleotides from the active site before the polymerase can incorporate the bases. Mismatches in the reads are mainly caused by spectral misassignments of the fluorophores used (Eid, et al., 2009). Deletions are likely caused by base incorporations that are faster than the rate of data recording (Eid, et al., 2009). The errors seem to be non-systematic, resulting in the lowest GC bias to compared to other platforms (Ross, et al., 2013).

The error rate of PB sequencing can be reduced through the use of circular consensus sequencing (CCS) (Travers, et al., 2010). In CCS, a hairpin adaptor is ligated to both sides of a linear DNA sequence. During sequencing, the polymerase can then pass multiple times over the same sequence (depending on the processivity of the polymerase). The multiple passes are be called into consensus and collapsed, yielding higher quality yet shorter reads, resulting in lower throughput. Consequently, many PB datasets generated do not utilize this methodology. Following this trend, the methods for overlap detection outlined in this paper have thus been designed for non-CCS reads.

### 2.2 Current challenges when using ONT sequencing

ONT sequencing works by measuring minute changes in ionic current across a membrane when a single DNA molecule is driven through a biological nanopore (Stoddart, et al., 2009). Currently, signal data is streamed to a cloud-based service called Metrichor that at the time of writing this paper, still uses hidden Markov models (HMM) with states for every possible 6-mer to render base calls on the data.

In the current HMM base calling methodology, if one state is identical to its next state, no net change in the sequence can be detected. This means that homopolymer states longer than five cannot be captured as they would be collapsed into a single 6-mer. It has also been observed that there are some 6-mers, particularly homopolymers, underrepresented in the data (Jain, et al., 2015; Loman, et al., 2015) when compared to the 6-mer content of the reference sequence, suggesting that there may be a systematic bias to transition in some states over others. In addition, there is some evidence suggesting GC biases within this type of data (Goodwin, et al., 2015; Laver, et al., 2015). We note that the base calling problem is under active development, with alternative base calling algorithms such as Nanocall (David, et al., 2016) and DeepNano (Boža, et al., 2016), recently made publicly available.

One can mitigate error rates in ONT data by generating two-direction (2D) reads. Similar to CCS for the PB platform, 2D sequencing involves ligating a hairpin adaptor, and allowing the nanopore to process both the forward and reverse strand of a sequence (Jain, et al., 2015). Combining information from both strands was shown to decrease the error rate from 40–30% to 10–20% with earlier chemistry (Jain, et al., 2015; Quick, et al., 2014), similar to the error rates of non-CCS PB sequencing. For the comparisons presented in this paper, we only consider 2D reads, as we expect investigators to prefer using higher quality ONT data.

## 3 Definitions and Concepts

In the context of DNA sequencing, an *overlap* is a broad term referring to a sequence match between two reads due to local regions on each read that originate from the same locus within a larger sequence (e.g., genome). The detail at which an overlap can be described can have large implications on both the downstream processing and computational costs associated with overlap computation, as discussed below.

Overlap between two reads may be *full* (complete) or *partial*, and may *dovetail* each other or one may be *contained* in the other (Fig. 1). The former pair of classifications often is a manifestation of data quality, but may also indicate haplotypic variations or polymorphisms. Full overlaps are overlaps that cover at least one end of a read in an overlap pair, whereas partial overlaps cover any portion of either read without the ends (Fig. 1). Sources contributing to observed partial overlaps include false positives due to near-repeats, chimeric sequences, or other artifacts (Li, 2016). Disambiguating the source of the overhang in partial overlaps may be important to downstream applications, especially when using non-haploid, metagenomic, and transcriptomic datasets.

**Fig. 1.**
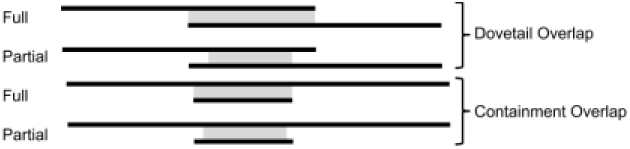
Visualization of partial and full overlaps in dovetail or containment forms.

The task of determining *overlap candidates* (Fig. 2A) is usually the first step in an overlap algorithm, and it refers to a simple binary pairing of properly oriented reads. To find overlap candidates on error-prone long reads, most methods look for matches of short sequence seeds (*k*mers) between the sequences. With a collection of overlap candidates, one can build a directionless overlap graph.

**Fig. 2.**
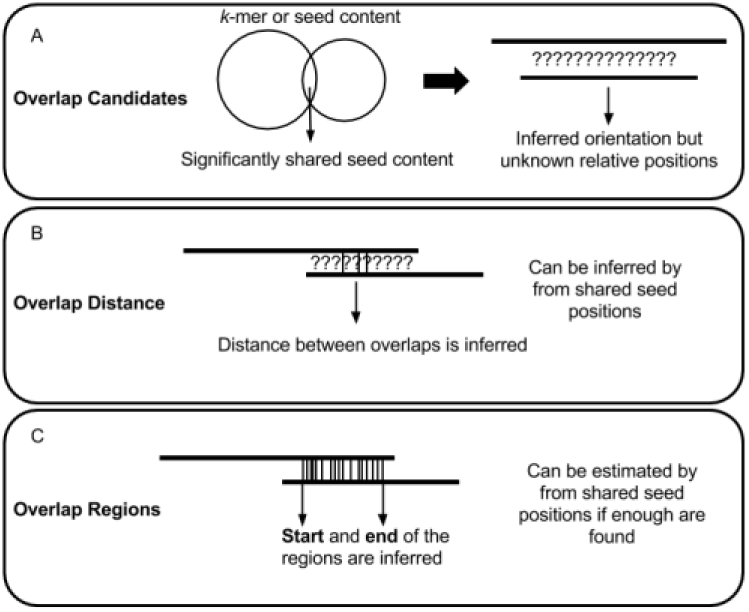
An overview of possible outcomes from an overlap detection algorithm. Each level has a computational cost associated with it, with the general trend being A<B<C. The common seeds-based comparison methods are not the only way to obtain these overlaps, but it is the most popular method used.

*Overlap distance* (Fig. 2B) refers to the relative positions between two overlapping reads. These distances provide directionality to the edges of the overlap graph. Theoretically, if the sequences are insertion or deletion (indel) -free, then a correct overlap distance would be sufficient to produce a layout and build a consensus from the reads. However, even a single indel error in one of the reads will cause a shift of coordinates, which would complicate consensus calling. Also, one cannot distinguish between partial and complete overlaps just with the distance information alone. Overlap distance can be estimated without a full alignment, based on a small number of shared seeds.

*Overlap regions* (Fig. 2C) refer to relative positions between overlapping reads, with the added information of start and end positions of the overlap along each read. If no errors are present, the sizes of the regions on both reads should be identical. In practice, due to high indel errors in long reads, this is rarely the case. Nevertheless, one can use this information to distinguish between partial and full overlaps. Similar to overlap distance, overlap regions can be estimated without a full alignment, but typically, more shared seeds are required for confident estimations.

There are many similarities between local alignments and overlaps. However, although a local aligner can serve as a read overlap tool (Chaisson and Tesler, 2012; Sović, et al., 2016), overlaps are not the same as local alignments. At the minimum, a read overlapper tool may simply indicate overlap candidates, rather than full base-to-base alignment coordinates. In some cases, similar to the discovery of partial overlaps, it may be important to find local alignments, as they may help discover repeats, chimeras, undetected vector sequences and other artifacts associated or confused with an overlap (Myers, 2014). Although, it is possible for an overlap detection algorithm to produce a full account of all the bases in overlapping reads, doing so would typically require costly algorithms like Smith-Waterman (Waterman, 1995).

All tools presented in this review have the ability to provide overlap candidates with at least an estimated local overlap region. Many tools can produce full local alignments, but due to the high computational costs, some tools provide an option for computing overlap regions and local alignments separately (such as GraphMap (Sović, et al., 2016)). Alternatively, other tools may not provide alignment refinement (such as MHAP (Berlin, et al., 2015) and Minimap (Li, 2016)). Indeed, since a full alignment between every pair of reads is not needed in some pipelines (Berlin, et al., 2015; Li, 2016), it is beneficial for overlap tools to be able to skip the computation of a full alignment.

## 4 Long Read Overlap Methodologies

Sequence overlap algorithms look for shared seeds between reads. Due to the high error rates of sequence reads on the PB and ONT platforms, these seeds tend to be very short (Supp. Figs. S1 and S2). The core differences between algorithms (Fig. 3) relate to not only how shared seeds are found, but in the way the seeds are used to determine an overlap candidate. After a method finds candidates, it will then refine them, usually using relative locations of the seeds within reads, and computing the estimated overlap regions. It is also common to check whether the candidate has a valid overlap by considering consistency of relative seed locations. Each method produces a list of overlap candidates, and provides an overlap region between reads. In some pipelines, the majority of the compute time is spent on realigning overlapping reads for error correction after candidates are found (Sović, et al., 2016). In others, precise alignments may not be needed (Li, 2016). Thus, the output of each overlap algorithm contains, at minimum, the overlap regions, and often with some auxiliary information for downstream applications (Table 1).

**Fig. 3.**
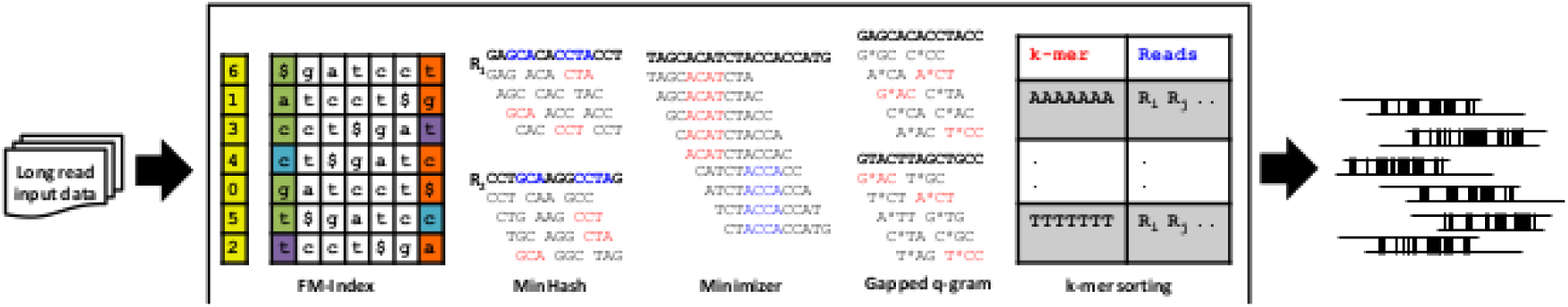
Visual overview of overlap detection algorithms. At the least, each method produces overlap regions. They may also generate auxiliary information, such as alignment trace points or full alignments.

**Table 1.** Summary of overlap tools output formats, associated pipelines, and availability.

| Software | Algorithm features | Associated assembly tools | Output | Availability |
| --- | --- | --- | --- | --- |
| BLASR | FM-Index, anchor clusters | PBcR | SAM alignment, other proprietary formats (overlap regions) | https://github.com/PacificBiosciences/blasr |
| Daligner | Cache efficient $k$-mer sort (radix) and merge | DAZZLER, MARVEL, FALCON | Local Alignments, LAS format (alignment tracepoints) | https://github.com/thegenemyers/DALIGNER |
| MHAP | MinHash | PBcR, Canu | MHAP output format (overlap regions) | https://github.com/marbl/MHAP |
| GraphMap | Gapped $q$-gram (spaced seeds), colinear clustering | Ra | SAM alignment, MHAP output format (overlap regions) | https://github.com/isovic/GraphMap |
| Minimap | Minimizer colinear clustering | Miniasm | PAF (overlap regions) | https://github.com/lh3/Minimap |

### 4.1 BLASR

BLASR was one of the first tools developed specifically to map PB data to a reference (Chaisson and Tesler, 2012). It utilizes methods adapted for fast short read alignments to long read data with high indel rates, combining concepts from Sanger-era and next-generation sequencing alignments. BLASR uses an FM-index (Ferragina, et al., 2005) to find short stretches of clustered alignment anchors (of length *k* or longer), generating a short list of candidate intervals/clusters to consider. A score is assigned to the clusters based on the frequency of alignment anchors. Top candidates are then processed into a full alignment.

Although BLASR was originally designed for read mapping, it has since been successfully used to produce overlaps for *de novo* assembly of several bacterial genomes (Chin, et al., 2013). However, to use the method for overlap detection one needs to carefully tune its parameters. For example, to achieve high sensitivity, BLASR needs prior knowledge of the read mapping frequency to parameterize nBest and nCandidates (default 10 for both) to a value higher than the coverage depth. Runtime of the tool is highly dependent on these two parameters (Berlin, et al., 2015), which may be due to the cost of computing a full alignment or the added computational cost per lookup to obtain more anchors.

What make this method slower are the choice of data structure, and its search for all possible candidates (not only the best candidates) for each lookup performed. The theoretical time complexity of a lookup in an FM-index data structure is linear with respect to the number of bases queried (Ferragina, et al., 2005), albeit not being very cache efficient (Myers, 2014). Thus, if one maps each read to a unique location, this would only take linear time with respect to the number of bases in the dataset. However, since only short (and often non-unique, cf. Supp. Figs. S1, S2) contiguous segments can be queried due to the high error rate, extra computation is required to consider all additional candidate anchor positions. Finally, BLASR computes full alignments rather than just overlap regions, thus, possibly utilizing more computational resources than needed for downstream processes.

### 4.2 DALIGNER

DALIGNER was the first tool designed specifically for finding read-to-read overlaps using PB data (Myers, 2014). This method focuses on optimizing the cache efficiency, in response to the relatively poor cache performance of the FM-index suffix array/tree data structure. It works by first splitting the reads into blocks, sorting the *k*-mers in each block, and then merging those blocks. The theoretical time complexity of DALIGNER is dominated by the merging step, which is quadratic in the number of occurrences of a given *k*-mer in a dataset (Myers, 2014).

To optimize speed and mitigate the effect of merging, DALIGNER filters out or decreases the occurrences of some *k*-mers in the dataset. Using a method called DUST (Morgulis, et al., 2006), DALIGNER (-mdust option) masks out low complexity regions (e.g. homopolymers) in the reads before the *k*-mers are extracted. Using a second method it filters out *k*-mers by multiplicity (-t option), increasing the speed of computation, decreasing memory usage, and mitigating the effects of repetitive sequences. However, these options also carry the risk of filtering out important *k*-mers needed for overlaps.

To use DALIGNER efficiently on larger datasets, splitting of the dataset into blocks is necessary. The comparisons required to perform all overlaps is quadratic in time relative to the number of blocks. Unique amongst the overlap tools, DALIGNER provides a means to split input data based on the total number of base pairs and read lengths (using the DBsplit utility). DALIGNER optionally outputs full overlaps, but will first output local alignment *tracepoints* to aid in computing a full alignment in later steps, producing large auxiliary files.

### 4.3 MHAP

MHAP (Berlin, et al., 2015) is a tool that uses the MinHash algorithm (Broder, 1997) to detect overlaps based on *k*-mer similarity between any two reads. MinHash computes the approximate similarity between two or more sets by hashing all the elements in the set with multiple hash functions, and storing the elements with the smallest hashed values (minimizers) in a *sketch* list. Using the minimum hash value is a form of locality-sensitive hashing, since it causes similar elements in a set to hash to the same value. In MHAP, overlap candidates are simply two *k*-mer sets that have a Jaccard index score above a predefined threshold.

After the overlap candidates are found, overlap regions are computed using the median relative positions of the shared minimizers. Because the sketch size used for each read is the same, MHAP may waste space and lose sensitivity if reads vary widely in length (Li, 2016). The time complexity of computing a single MinHash sketch is O(*kl*), where *l* is the number of *k*-mers in the read set. Thus, evaluating *n* reads for a sketch size *h* for all resemblances takes O((*hn*)^2^) time (Broder, 1997). MHAP further reduces its time complexity by storing *h* min-mers in *h* hash tables to use for lookups to find similar reads (Berlin, et al., 2015).

Like DALIGNER, MHAP functions best when repetitive elements are not used as seeds. MHAP supports the input of a list of *k*-mers, ordered by multiplicity, obtained by using a 3rd party *k*-mer counting tool, such as Jellyfish (Marçais and Kingsford, 2011). We note that, although this strategy would improve the performance of the tool, the computational cost of *k*-mer counting may not be trivial.

MHAP’s computational performance is confounded by its implementation. While most high-performance bioinformatics tools utilize C/C++ for their performance benefits, MHAP is implemented in Java. Another method called Minlookup (Wang and Jones, 2015), written in C, utilizes a similar algorithm to MHAP, however it is designed with ONT datasets in mind. The authors demonstrate improved performance associated with their implementation. However, Minlookup was not evaluated here as it is in early development, and cannot use multiple CPU threads.

### 4.4 GraphMap

GraphMap, like BLASR, was designed primarily as a read mapping tool (Sović, et al., 2016), but for ONT data. It specifically addresses the overlap detection problem, notably producing full alignments. GraphMap also provides an option to generate overlap regions exclusively.

In GraphMap the “-owler” option activates a mode specifically designed for computing overlaps. Like its standard mapping algorithm, it first creates a hash table of seeds from the entire dataset. The seeds it uses are not *k*-mers, but rather gapped q-grams (Burkhardt, et al., 2002) – *k*-mers with wild card positions, also called spaced seeds (Keich, et al., 2004). It is not clear what gapped q-grams work optimally with ONT or PB data; more research is needed to determine the optimal seeds to cope with high error rates. The current implementation uses a hardcoded seed that is 12 bases long with an indel/mismatch allowed in the middle (6 matching bases, 1 indel/mismatch base, followed by 6 matching bases). GraphMap then collects seed hits, using them for finding the longest common subsequence in k-length substrings (Benson et al., 2013). The output from this step is then filtered to find collinear chains of seeds (private correspondence with Ivan Sović). The bounds of these chains are then returned, using the MHAP output format.

### 4.5 Minimap

Minimap (Li, 2016) is an overlapper/mapping tool that combines concepts from many of its predecessors, such as DALIGNER (*k*-mer sorting for cache efficiency), MHAP (computing minimizers) and GraphMap (clustering collinear chains of matching seeds). Minimap subsamples the hashed *k*-mer space by computing minimizers, and compiles the corresponding *k*-mers along with their location on their originating reads.

Like MHAP, the use of repetitive *k*-mers as the min-*k*-mer can degrade the performance of overlap detection. To minimize the effect of repetitive elements, Minimap uses an invertible hash function when choosing min-*k*-mers. This is similar to DALIGNER’s use of DUST; it works by preventing certain hash values that correspond to low complexity sequences.

Also similar to DALIGNER, Minimap was designed with cache efficiency in mind. It stores its lists of minimizers initially in an array, which is later sorted for the seed merging step. Though the computational cost incurred by sorting the list can negatively impact performance compared with the constant cost of insertion in a hash table, its cache performance outperforms a conventional hash table. All hits between two reads are then collected using this sorted set, and are clustered together into approximately collinear hits. The overlap regions for each pair of overlaps are then finally outputted in pairing mapping format (PAF) (Li, 2016).

## 5 Benchmarking

We profiled and compared results from BLASR, DALIGNER, MHAP, GraphMap, and Minimap, using publicly available long read datasets with the newest chemistries available at the time of the study (Supp. Table S1). For PB we used *E. coli* (P6-C4) and *C. elegans* whole genome shotgun sequencing datasets. For ONT we used an *E. coli* (SQK-MAP-006) dataset. We also used simulated *E. coli* datasets for the PB and ONT platforms using PBSim (Ono et al., 2013) and NanoSim (Yang, et al., 2016), respectively, and simulated ONT *C. elegans* reads using NanoSim (Supp. Table S1). Only the *E. coli* datasets were used in a parameter sweep for in-depth evaluations of performance.

### 5.1 Sensitivity and FDR

We profiled the sensitivity and false discovery rate (FDR = 1 − precision) on the experimental PB P6-C4 *E. coli* and the ONT SQK-MAP-006 *E. coli* datasets. We also evaluated the tools on simulated data generated based on these datasets. Our ground truth for the real dataset was determined via bwa mem alignments to a reference, using −x pacbio and ont2d options, respectively (Li and Durbin, 2009). We note that these align ments may have missing or false alignments. However, these can still serve as a good estimate for ground truth comparisons, since mismatch rate to a reference is much lower than the observed mismatch between overlapping reads. In the latter case, reads that are, say, 80% accurate will have a mutual agreement of 64% on average. In addition, due to our reference-based approach, our metrics are resilient against false overlaps caused by repetitive elements. Further, all tools are compared against the same alignments; hence we expect our analysis to preserve the relative performance of tools. Finally, there is no ambiguity for ground truth in the simulated datasets, as each simulation tool reports exactly where in the genome the reads were derived from, allowing us to calculate the exact precision and sensitivity of each method.

To produce a fair comparison of the tools, we used a variety of parameters for each (Supp. note S1). These parameters were chosen based on tool documentation, personal correspondence with the tool authors, as well as our current understanding of their algorithms. We ran MHAP with a list of *k*-mer counts derived from Jellyfish (Marçais and Kingsford, 2011) for each value of *k* tested to help filter repetitive *k*-mers. GraphMap could not be parameterized when running in the “owler” mode, and had only one set of running parameters.

We counted an overlap as correct when the overlapping pair was present in our ground truth with the correct strand orientation. We did not take into account reported lengths of overlap, but note that this information may be important (e.g. to improve performance of realignment). For each tool we computed the skyline, or Pareto-optimal results, in our tests (the points with the highest sensitivity for a given FDR), and plotted these results on receiver operating characteristic (ROC)-like plots (featuring FDR rather than the traditional false positive rate).

We can see that although many tools have similar sensitivity and FDR depending on the parameterization, the overall trends reveal differences in sensitivity and FDR on each specific datatype (Fig. 4). For instance, MHAP can achieve high sensitivity on all datasets but lacks precision compared to most other methods on the ONT datasets. The only other tool that may have less precision on the ONT datasets is BLASR. DALIGNER proves to have a high sensitivity and precision, but it is not always the winner, especially on the ONT dataset. Minimap has high sensitivity and precision on the ONT datasets but does not maintain such performance on the PB dataset. Finally, the results for GraphMap were competitive despite using a single parameterization.

**Fig. 4.**
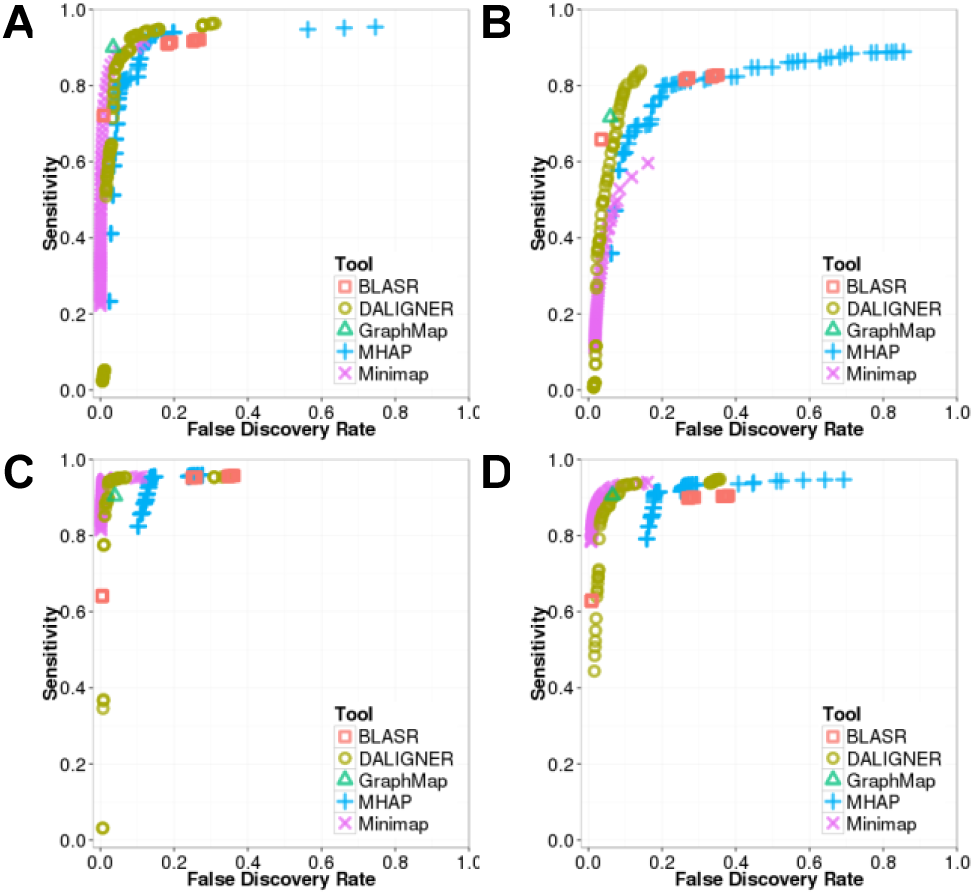
ROC-like plot on BLASR, DALIGNER, GraphMap, MHAP, GraphMap, and MHAP. A: PB *E. coli* simulated with PBsim. B: PB P6-C4 *E. coli* dataset. C: ONT *E. coli* simulated with Nanosim. D: ONT SQK-MAP-006 *E. coli* dataset.

These plots reveal that selection of operating parameters very much depends on the balance of project-specific importance attributed to sensitivity and precision, as expected. For instance, the importance of sensitivity is clear as it provides critical starting material for downstream processing. On the other hand, low sensitivity can be tolerated if the downstream method employs multiple iterations of error correction, because as errors are resolved within each iteration, the sensitivity is expected to increase. However, these downstream operations of course may come with a high computing cost.

The F1 score (also F-score or F-measure) represents a common way to combine these two score. It is the harmonic mean between the sensitivity and precision. To better compare these methods, we computed F1 scores for each using a range of parameters, and considered the highest value for each method to be representative of its overall performance. We calculated confidence intervals for the F1 scores using three standard deviations around the observed values, which revealed that reported F1 values were statistically significantly different from each other.

For the simulated PB data, GraphMap has the highest F1 score (despite being designed for ONT data and not PB data) followed by DALIGNER, Minimap, MHAP, and BLASR (Table 2). For the real PB data DALIGNER has the highest F1 score followed by GraphMap, MHAP, and BLASR. For both the simulated and real ONT datasets, Minimap was the best method, yielding the highest F1 score, followed by GraphMap, MHAP and BLASR (Table 2).

**Table 2.** An overview of sensitivity and precision on simulated and real error-prone long read datasets. In both the PB and ONT simulated datasets, the best values, shown in **bold** face, are statistically significantly better than the other values.

| Tool | Simulated PB *E. coli* |  |  | Simulated ONT *E. coli* |  |  | PB P6-C4 *E. coli* |  |  | ONT SQK-MAP-006 *E. coli* |  |  |
| --- | --- | --- | --- | --- | --- | --- | --- | --- | --- | --- | --- | --- |
|  | Sens. (%) | Prec. (%) | F1 (%) | Sens. (%) | Prec. (%) | F1 (%) | Sens. (%) | Prec. (%) | F1 (%) | Sens. (%) | Prec. (%) | F1 (%) |
| BLASR | 91.0 | 81.9 | 86.2 | **95.2** | 75.1 | 84.0 | 66.0 | **96.5** | 78.3 | 89.9 | 73.0 | 80.6 |
| DALIGNER | **92.4** | 91.9 | 92.1 | 94.9 | 97.6 | 95.9 | **83.8** | 85.8 | **84.8** | **92.9** | 91.0 | 91.9 |
| MHAP | 91.5 | 88.0 | 89.8 | **95.1** | 86.5 | 90.6 | 79.8 | 79.8 | 79.8 | 91.2 | 82.0 | 86.3 |
| GraphMap | 90.1 | **96.5** | **93.1** | 90.4 | 96.0 | 93.1 | 71.7 | 94.0 | 81.4 | 90.6 | 93.4 | 92.0 |
| Minimap | 88.9 | 94.8 | 91.8 | 94.6 | **99.0** | **96.7** | 59.6 | 83.8 | 69.7 | 91.2 | **95.4** | **93.2** |

Overall, these results suggest that some tools may perform substantially differently on data from different platforms. We hypothesize that, differences in the read length distributions and error type frequencies, could be responsible for this behaviour.

### 5.2 Computational Performance

To measure the computational performance of each method, we ran each tool with default parameters (with some exceptions see Supp. note S2) as well as another run with optimized parameters yielding the highest F1 score (Table 2 and Supp. note S1) obtained after a parameter sweep on the simulated datasets. We note that GraphMap’s owler mode could not be parameterized, except for choosing the number of threads, so there was no difference in the settings for default and highest F1 score parameterization runs. We ran our tests serially on the same 64-core Intel Xeon CPU E7-8867 v3 @ 2.50GHz machine with 2.5TB of memory. We measured the peak memory, CPU and wall clock time across read subsets to show the scalability of each method.

We investigated the scalability of each method, testing each method using 4, 8, 16 or 32 threads of execution on the *E. coli* datasets (Supp. Figs. S3–10). Despite specifying the number of threads, each tool often used more or fewer threads than expected (Supp. Figs. S3, S4, S11, S12). In particular, MHAP tended to use more threads than the number we specified.

On all tested *E. coli* datasets in our study, we observe that Minimap is the most computationally efficient tool, robustly producing overlap regions at least 3–4 times faster than all other methods, even when parameterizing for optimal F1 score (Supp. Fig. S7, S8). Determining the next fastest method is confounded by the effect of parameterization. For instance when considering only our F1 score optimized settings, DALIGNER, generally keeping within an order of magnitude or less of Minimap. On the other hand, DALIGNER can be 2–5 times slower than MHAP on some datasets under default parameters.

With default settings, DALIGNER performs up to 10 times slower than F1 score optimized settings. This primarily occurs because the *k*-mer filtering threshold (−t) in the F1 optimized parameterization not only increases specificity but also reduces runtime. In contrast, our parameterization to optimize the F1 score in MHAP decreases the speed (by a factor of 3–4). In this case, the culprit was the sketch size (--num-hashes) used; larger sketch sizes increase sensitivity at the cost of time.

Finally, GraphMap is generally the least scalable method, the slowest when considering default parameters only, and only 1–2 times faster than BLASR when considering F1 optimized settings. BLASR is also able to scale better than GraphMap to a more threads (Supp. Figs. S7, S8).

In addition to its impressive computational performance, Minimap uses less memory than almost all methods on tested *E. coli* datasets (Supp. Figs S9, S10), staying within an order of magnitude of BLASR on average, despite the latter employing an FM-index. Memory usage in GraphMap seems to scale linearly with the number of reads at a rate nearly 10 times that of the BLASR or Minimap, likely owing to the hash table it uses. The memory usage characteristics of DALIGNER and MHAP are less clear, drastically changing given the parameters utilized. Overall MHAP has the worst memory performance even when using default parameters. The cause of the memory increase between optimized F1 and default setting in MHAP is again due to an increase in the sketch size between runs. Because of *k*-mer multiplicity filtering, DALIGNER’s memory usage is 2–3 times lower when parameterized for an optimized F1 score.

Many of the trends from the *C. elegans* datasets mirror the performance on the smaller *E. coli* dataset. Again, computational performance on the larger *C. elegans* datasets is still dominated by Minimap (Supp. Figs. S11, S12), being at least 5 times faster than any other method. DALIGNER’s performance seems to generally scale well, especially when *k*-mer filtering is performed (within an order of magnitude of Minimap). With default settings, MHAP is 2–3 times faster than DALIGNER, but is several orders of magnitude slower, when the F1 score is optimized. The performance of GraphMap shows that it does not scale well to large number of reads (>100000), and its calculations take an order of magnitude longer than BLASR.

The performance of BLASR is a bit perplexing. When we use no *k*-mer filtering in DALIGNER, BLASR is within an order of magnitude of speed. When using optimized parameters, BLASR is also at least twice as fast as MHAP, which becomes more evident when using a large number (>15) of threads. We note that these results seemingly contradict the results found in previous studies (Berlin, et al., 2015; Myers, 2014). This may be due to different datasets and technology versions used by the two studies. It may also highlight the effects of parameterization of each tool – a difficult but critical task when tuning the performance of these tools.

The trends in memory performance on the *C. elegans* datasets are generally consistent with *E*. *coli* datasets (Supp. Fig. S11, S12). A notable exception however is the memory usage of DALIGNER, which begins leveling off with increased number of reads. Unlike with the *E. coli* dataset, this dataset is large enough that DALIGNER begins to split the data into batches, reducing its memory usage.

## 6 Discussion

Our study highlights that there are important considerations to factor in while developing new or improving existing tools.

### 6.1 Modularity

A tool that can report intermediate results may help reduce computation in downstream applications. For example, modularizing overlap candidate detection, overlap validation, and alignment can provide flexibility when used in different pipelines. Graphmap’s owler mode is an example of this, enabling users to generate MHAP-like output for overlap regions, rather than a more detailed alignment on detected regions. Further, compliance to standardized output is highly recommended, including for generating intermediate results. Doing so would not only allow one to perform comparative performance evaluations on a variety of equivalent metrics, but also allow for flexibility in creating new pipelines. Examples of emergent output standards include the Graphical Fragment Assembly (GFA) (https://github.com/pmelsted/GFA-spec) format, PAF (Li, 2016), and the MHAP output format.

### 6.2 Cache Efficiency

Given the concepts presented, and along with our benchmarks performed herein indicates that theoretical performance estimations based on time complexity analysis might not be enough to conclude on what works best. Traditional algorithm complexity analysis suffers from an assumption that all memory access costs are the same. However, on modern computers intermediate levels of fast-access cache exist between the registers of the CPU and main memory. A failed attempt to read or write data in the cache is called a cache miss, causing delays by requiring the algorithm to fetch data from other cache levels or main memory.

Cache efficiency in algorithmic design has become a major consideration, and in some cases will trump many time complexity based motiva tions for algorithmic development. For instance, though the expected time complexity of DALIGNER has a quadratic component based on the number of occurrences of a *k*-mer in the dataset, its actual computational performance seems to be much better empirically. The authors claim this is due to the cache efficiency of the method (compared to using an FM-index) (Myers, 2014), and in practice this also seems to be the case, as observed in our comparisons.

The basic concept of a cache efficient algorithm relies on minimizing random access whenever possible, by serializing data accesses in blocks that are small enough to fit into various levels of cache, especially at the levels of cache with the lowest latency. Algorithms that exploit a specific cache configuration utilize an I/O-model (also called the external-memory model) (Aggarwal, et al., 1988; Demaine, 2002). Conceptually, these algorithms must have explicit knowledge of the size of each component of the memory hierarchy, and will adjust the size of contiguous blocks of data to minimize data transfers from memory to cache.

In contrast to the I/O model, algorithms that are designed with cache in mind, but do not explicitly rely on known cache size blocks are called cache oblivious (Frigo, et al., 1999). Cache oblivious algorithms are beneficial, as they do not rely on the knowledge of the processor architecture; instead they utilize classes of algorithms that are inherently cache efficient such as scanning algorithms (e.g. DALIGNER’s merging step of a sorted list).

### 6.3 Batching and Batch/Block Sizes

For many of the methods surveyed in this paper, memory usage can be roughly quadratic relative to the number of reads, and at least linear to the number of *k*-mers in the set. Thus, to perform all necessary comparisons (i.e. to compute an upper triangular matrix of candidate comparisons), the data must be processed in batches. Generally, it is better to use as few blocks as possible, since the time required to perform all overlaps is quadratic relative to the number of batches. Methods that have a very low memory usage overall will be able to have the computational benefit of splitting the data into fewer batches. Batching is handled in different ways depending on the tool. Some tools have built-in splitting (DALIGNER/DAZZLER database with DBsplit), and others have this process built into their associated pipelines (e.g. MHAP and PBcR). Other methods (BLASR, Minimap) seem to have more scalable memory requirements, and may not require splitting.

### 6.4 Repetitive elements and sequence filtering

Any common regions due to homology or other repetitive elements may confound read-to-read overlaps, and may be difficult to disambiguate from true overlaps. Such repetitive elements may lead to many false positives in overlap detection, and may increase the computational burden, leading to lower quality in downstream assembly. Thus, it is common for overlap methods to employ sequence filtering, by removal or masking of repetitive elements to improve algorithmic performance both in run time and specificity. Many of the methods compared utilize *k*-mer frequencies to filter highly repetitive *k*-mers using an absolute or percent *k*-mer multiplicity. Another common filtering strategy is to prevent the use of low complexity sequences.

## 7 Conclusions

There are many challenges in evaluating algorithms that function on error-prone long reads, such as those from PB and ONT instruments. Although both sequencing technologies have comparable error rates, characteristics of their errors as well as their read length distributions are substantially different. Also, within each technology there are rapid improvements in quality (Jain, et al., 2015; Laver, et al., 2015), causing disagreement between datasets derived from the same technology.

Despite these issues, we show that Minimap is the most computationally efficient method (in both time and memory) and is the most specific and sensitive method on the ONT datasets tested. We note that Minimap is not as sensitive or as specific as Graphmap, DALIGNER or MHAP on the PB datasets tested. Our results shown that GraphMap and DALIGNER are most specific and sensitive method on PB datasets tested, though DALIGNER scales better computationally. PB being a more mature technology compared to ONT, it is not surprising to see several tools performing well on the platform.

Here, we have provided an overview of read-to-read overlap detection concepts, comparing leading methods for researchers can make informed decisions given their datasets and computational resources. We hope that our elucidation to open problems and key concepts to consider will be a helpful resource for those looking to develop new or improve on existing overlap detection tools.

## Acknowledgements

The authors would like to thank Sergey Koren, Ivan Sović for their help and suggestions when running MHAP and GraphMap, respectively, as well as their insights into the behaviour and results of each tool on different datasets.

